# RNA-protein binding motifs mining with a new hybrid deep learning based cross-domain knowledge integration approach

**DOI:** 10.1101/085191

**Authors:** Xiaoyong Pan, Hong-Bin Shen

**Affiliations:** Department of Veterinary Clinical and Animal Sciences, University of Copenhagen, Copenhagen, Denmark; Institute of Image Processing and Pattern Recognition, Shanghai Jiao Tong University, and Key Laboratory of System Control and Information Processing, Ministry of Education of China, Shanghai, China

**Author notes:** Shared Correspondence.

**Keywords:** RNA-binding protein, CLIP-seq, deep belief network, convolutional neural network, multimodal deep learning

## Abstract

**Background:** RNAs play key roles in cells through the interactions with proteins known as the RNA-binding proteins (RBP) and their binding motifs enable crucial understanding of the post-transcriptional regulation of RNAs. How the RBPs correctly recognize the target RNAs and why they bind specific positions is still far from clear. Machine learning-based algorithms are widely acknowledged to be capable of speeding up this process. Although many automatic tools have been developed to predict the RNA-protein binding sites from the rapidly growing multi-resource data, e.g. sequence, structure, their domain specific features and formats have posed significant computational challenges. One of current difficulties is that the cross-source shared common knowledge is at a higher abstraction level beyond the observed data, resulting in a low efficiency of direct integration of observed data across domains. The other difficulty is how to interpret the prediction results. Existing approaches tend to terminate after outputting the potential discrete binding sites on the sequences, but how to assemble them into the meaningful binding motifs is a topic worth of further investigation.

**Results:** In viewing of these challenges, we propose a deep learning-based framework (iDeep) by using a novel hybrid convolutional neural network and deep belief network to predict the RBP interaction sites and motifs on RNAs. This new protocol is featured by transforming the original observed data into a high-level abstraction feature space using multiple layers of learning blocks, where the shared representations across different domains are integrated. To validate our iDeep method, we performed experiments on 31 large-scale CLIP-seq datasets, and our results show that by integrating multiple sources of data, the average AUC can be improved by 8% compared to the best single-source-based predictor; and through cross-domain knowledge integration at an abstraction level, it outperforms the state-of-the-art predictors by 6%. Besides the overall enhanced prediction performance, the convolutional neural network module embedded in iDeep is also able to automatically capture the interpretable binding motifs for RBPs. Large-scale experiments demonstrate that these mined binding motifs agree well with the experimentally verified results, suggesting iDeep is a promising approach in the real-world applications.

**Conclusion:** The iDeep framework not only can achieve promising performance than the state-of-the-art predictors, but also easily capture interpretable binding motifs. iDeep is available at http://www.csbio.sjtu.edu.cn/bioinf/iDeep

## Background

RNA-protein interactions are involved in many biological processes, such as gene regulation and splicing [1]. Discovering the RNA-protein interactions has a great potential for further understanding the mechanisms behind those biological processes. For example, Argonaute (AGO) protein belongs to components of the RNA-induced silencing complex (RISC), which transfers microRNAs (miRNAs) to be bound with their target genes, thereby inhibit target gene expression [2]. Sequence-specific associations between RBPs and their RNA targets are mediated by binding domains, which recognize binding sites on RNAs. Where the RNA-protein binding sites on the RNAs are usually short sequences with 4 to 30 base pairs long, typically referred as binding motif. Detecting them can facilitate the deeper insights into post-transcriptional regulation.

Although there are many genome-wide RNA-binding protein detection techniques, such as RNAcompete [3], PAR-CLIP [4], they are still cost-heavy and time-intensive. Fortunately, with the advent of these high-throughput techniques, many useful genome-wide data associated with RBPs are generated rapidly, including specific binding positions on RNAs with proteins. These data provides important bases for developing computational approaches to predict the RBP binding sites by using the advanced computational methods [5, 6, 7, 8, 9].

At the very beginning of the methodology development of this field, predictors are mainly constructed by only using the sequence information. For instance, MatrixREDUCE simply fits a statistical mechanical model to infer the sequence-specific binding sites for transcription factors from sequences [10]. DRIMust discovers motifs by integrating the minimum hyper-geometric statistical framework with suffix trees for fast enumerating motifs [11].

Besides the high-throughput sequences, actually multiple sources of data are available from the genome-wide RNA-protein CLIP-seq data, such as sequences, structures, genomic context. Each source of data has a different kind of representation and correlation structure. A popular straightforward idea is to integrate these data to construct a predictor, which is expected to be very useful for enhancing the prediction accuracy. Two integration schemes have been widely used in the literatures:

1. Feature-level fusion. This type of fusion strategy is to encode the different sources into feature vectors, which will be concatenated together. For instance, the OliMoSS model has integrated tetranucleotide sequence, binding motifs and secondary structures to predict protein specific interactions on RNAs by simply concatenating the different sources of features into one high-dimensional features (525-D) [12], which may result in difficulties for the following statistical learning process. For instance, the learning algorithm used in the OliMoSS is support vector machine (SVM), which will easily suffer from the curse of dimension problem. Similar strategy is also applied in DNA-protein binding sites prediction [13]. The other implementation of feature-level fusion is the multiple-kernel learning, which design multiple kernels for different features, and then combine them together [14, 15]. Similarly, GraphProt encodes the sequence and structure information to graph kernel to predict binding reference of RBPs [6].
2. Decision-level fusion. To solve the high-dimension space learning problem, decision level-based fusion system has been proposed. For instance, the iONMF [5] is a predictor for predicting RNA-protein interaction sites. It has trained a model for each of available resource data, e.g. kmer sequence, secondary structure, CLIP co-binding, Gene Ontology (GO) information, and region type. These independent 5 models will work independently, which have no interconnections between them during the training processes. The final prediction output of the whole system is the fusion of 5 independent predictions.

Despite the progresses of previously proposed methods, they have a shared drawback that the models were constructed on the features extracted from the observed data, where the frequent noise may make the subsequent classifiers learn wrong knowledge. Deep learning [16, 17] is a recently developed approach, which works in a hybrid multiple-layer abstraction way by mapping the observed data to a much high-level abstraction space, where the prediction model will be constructed. This new type of approach has provided much attractive solutions for integrating heterogeneous data and are effective in automatically learning complex patterns from multiple simple raw inputs.

One typical deep learning framework is known as the convolutional neural network (CNN) [18]. The advantage of CNN is that it does not separate feature extraction and model learning into two independent steps any more as done in the traditional statistical learning algorithms. Instead it simultaneously learns features and classification models from the original input in a data-driven way, which will reduce the potential mismatch effects between the feature extraction and learning classification models. The CNN model has been applied in the binding proteins prediction of DNA or RNA. For instance, a recent CNN-based deep learning approach Deep-Bind was proposed to predict sequence specificities for protein binding RNA/DNA [8]. Similarly, the DeepSEA [19] utilizes the deep CNNs to learn regulatory sequence motifs for predicting DNA functions from chromatin profiling data; Basset [20] trained analogous deep CNN models to learn impacts of DNA sequences variants on chromatin regulation from large-scale DNase-seq data. These studies have shown that the convolution operation in CNN is able to scan a set of weight matrix (filters) across the input sequences to recognize relevant patterns that respond to motifs, like patterns corresponding to edges and curve fragments in images [21, 22], resulting in better prediction accuracies [12, 5].

The deep belief network (DBN) is another deep learning algorithm to learn high-level features from large-scale data [26], which is also a recent popular choice for constructing the computational models. For example, the deepnet-rbp fused the structural and k-mer sequence features to predict RBP interaction sites [23] using DBNs. DANN trains a DBN to annotate non-coding variants [24], which is able to capture non-linear abstraction features. We also developed a model called IPMiner by applying the stacked autoencoder to learn high-level features for predicting RNA-protein interactions from raw sequence composition features, and it yielded promising performance compared to other sequence-based methods [25]. It’s worth noting that many studies have shown that the CNN and DBN hold their own advantages due to different deep learning architectures, e.g. CNN is more appropriate for sequence data and DBN prefers the numeric inputs. This motivates us to consider how to integrate the merits of CNN and DBN for better prediction of RBP binding sites and find the sequence motifs.

In this study, we propose a multimodal deep learning framework iDeep, a hybrid framework with CNNs and DBNs, to better integrate multiple heterogeneous data sources for predicting RBP interaction sites on RNAs (Fig. 1). For the data represented by the binary or numeric features, the DBN networks will be used; While for the sequence data, the CNN network will be applied. Different deep network models will be trained and tuned together from the top shared layer to the individual bottom layers using backpropagation, and then the shared latent features are captured across them. Compared to the existing approaches, the iDeep has the following merits: 1) the iDeep is constructed with a deep learning structure, and it consists of multiple neural networks stacked together [17, 16], where the outputs of each layer are the inputs of successive layer. Such layer-by-layer learning helps to reduce the noise effects in the original input. 2) The iDeep successfully integrates the CNN and DBN for dealing with the different sources of protein-RNA binding related data to enhance the discrimination ability. The CNN is able to capture regulatory motifs, which are recurring patterns in RNA sequences with a biological function. The DBN learns high-level features regarded as a joint distribution determined by hidden variables for different inputs. 3) The hybrid framework of flexible multimodal learning and fusion at an abstraction level makes the iDeep handle different features in an easy manner. The top shared hidden layer at the fusion level will help discover the shared properties across different modalities [27, 28].

**Figure 1.**
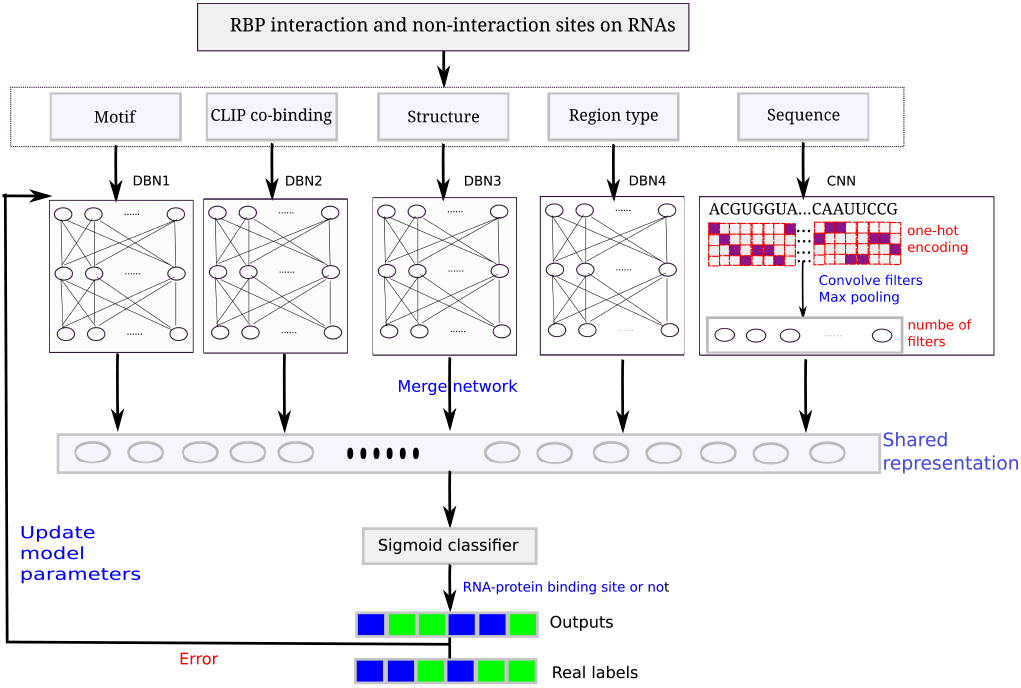
**The flowchart of proposed iDeep for predicting RNA-protein binding sites on RNAs.** It firstly extracted different representation for RNA-protein binding sites within a windows size 101, then use multimodal deep learning consisting of DBNs and CNNs to integrate these extracted representations to predict RBP interaction sites.

## Results

In this study, we evaluated iDeep on independent testing datasets, and also compared it with the performance of DBN and CNN from individual sources of data. To demonstrate the advantage of iDeep, some state-of-art predictors of iONMF, DeepBind, and Oli were also compared. Besides, a large-scale analysis has been conducted to demonstrate the discovered binding motifs using iDeep.

### The iDeep’s performance

To demonstrate the ability of iDeep for predicting RNA-protein binding sites, we evaluate iDeep on independent testing dataset (see the dataset section). We firstly use 4000 training samples for model training, 1000 validation sites are evaluated at the end of each training epoch to monitor the convergence. For each experiment, iDeep is trained with the same initializations. After we obtain the trained model, we apply it to predict binding sites for 1000 independent testing samples. The ROC on 31 experiments are shown in Fig. 2. It indicates that iDeep yields different performance on different experiments with huge margin, the AUC ranges from 0.70 for protein hnRNPL1-like to 0.98 for protein PUM2. In addition, iDeep achieves the AUC greater than 0.90 on 23 of 31 experiments, and the average AUC of iDeep on all experiments is 0.90, indicating that iDeep accurately predict RBP binding sites on a genome-wide scale.

**Figure 2.**
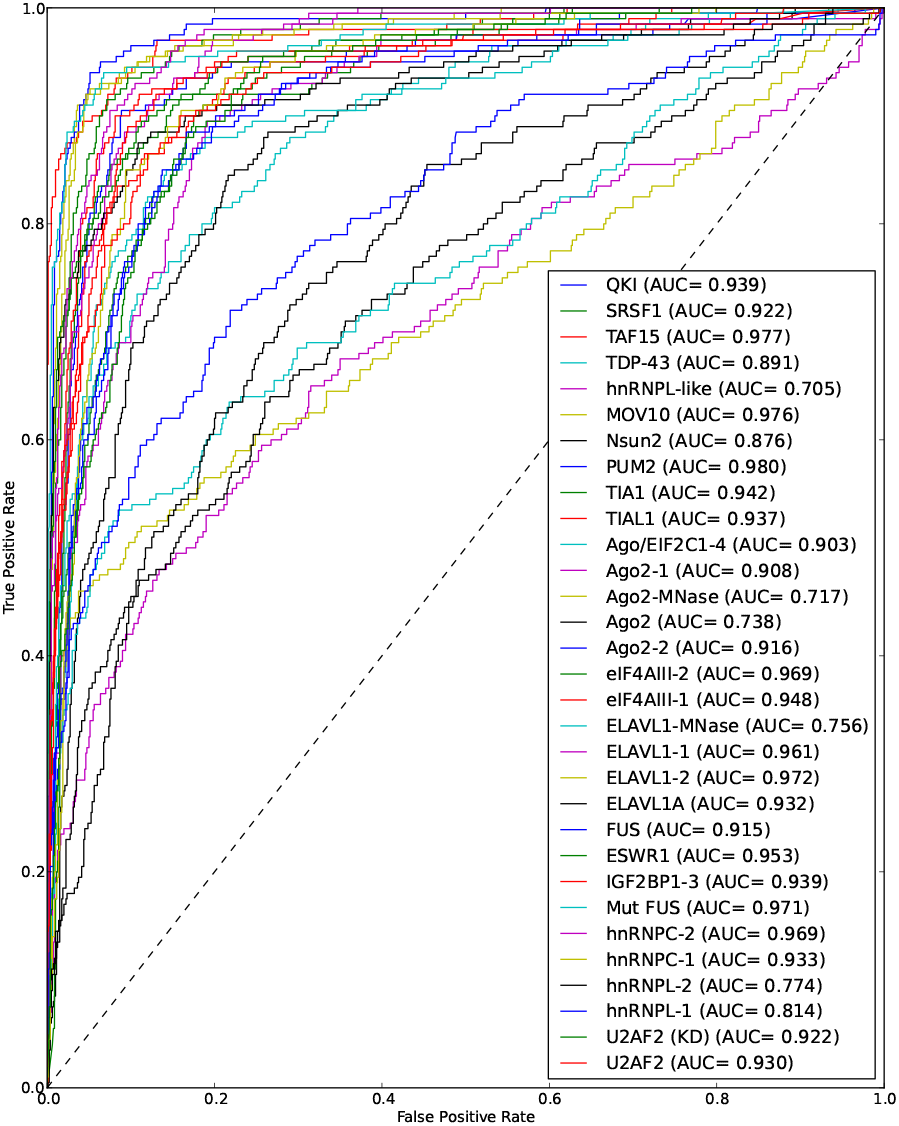
**ROC Performance.** The ROC curve for predicting RNA-protein binding sites on 31 experiment dataset.

### Comparing iDeep with other state-of-the-art methods

We firstly compare it with state-of-the-art method iONMF, which has shown better performance than other existing methods [5], such as GraphProt [6] and RNAContext [37]. As shown in Table 2, we can see that iDeep outperform iONMF on most of the 31 experiments, the average AUC of the 31 experiments increases from 0.85±0.08 of iONMF to 0.90±0.09 of iDeep. Furthermore, for some experiments, it improves the AUC over 15%, such as for protein hnRNPL-2, the AUC increases from 0.66 of iONMF to 0.77 of iDeep. In addition, iDeep also performes better than other matrix factorization-based methods NMF [38], SNMF [39] and QNO [40], which achieves the average AUC of 0.83±0.10, 0.71±0.14, 0.79±0.12 on 31 experiments, respectively.

We further compare iDeep with another protein-specific method Oli [12], which yields an average AUC of 0.77±0.16, and 17% lower than the iDeep. We find that it has a bigger performance variance than other tested methods. For example, Oli performs very bad on some experiments, e.g. AUC 0.39 on hnRNPL-1 protein, but on some experiments, its performance is very good, e.g. 0.94 on PUM2 protein. For the DeepBind [8] approach using the same parameters of CNN integrated in iDeep, it achieves an average AUC 0.83±0.12 across 31 experiments, which performs worse than iDeep. The reason is that DeepBind cannot yield promising performance across all 31 experiments from only sequences.

To demonstrate the merits of the designed framework of iDeep, we also compare iDeep with its own variant iDeep-kmer, whose input modalities are kmer, region type, clip-cobinding and structure using the same network architecture. The only difference is that iDeep uses CNN sequence and motif modalities instead of high-dimensional kmer modality. As indicated in Table 1, iDeep-kmer yields an average AUC of 0.87±0.09, which is worse than iDeep, indicating that CNN sequence and motif modality have better discriminating ability than high-dimensional kmer modality. On the other hand, iDeep performs faster than iDeep-kmer both in the training and testing steps.

**Table 1.**
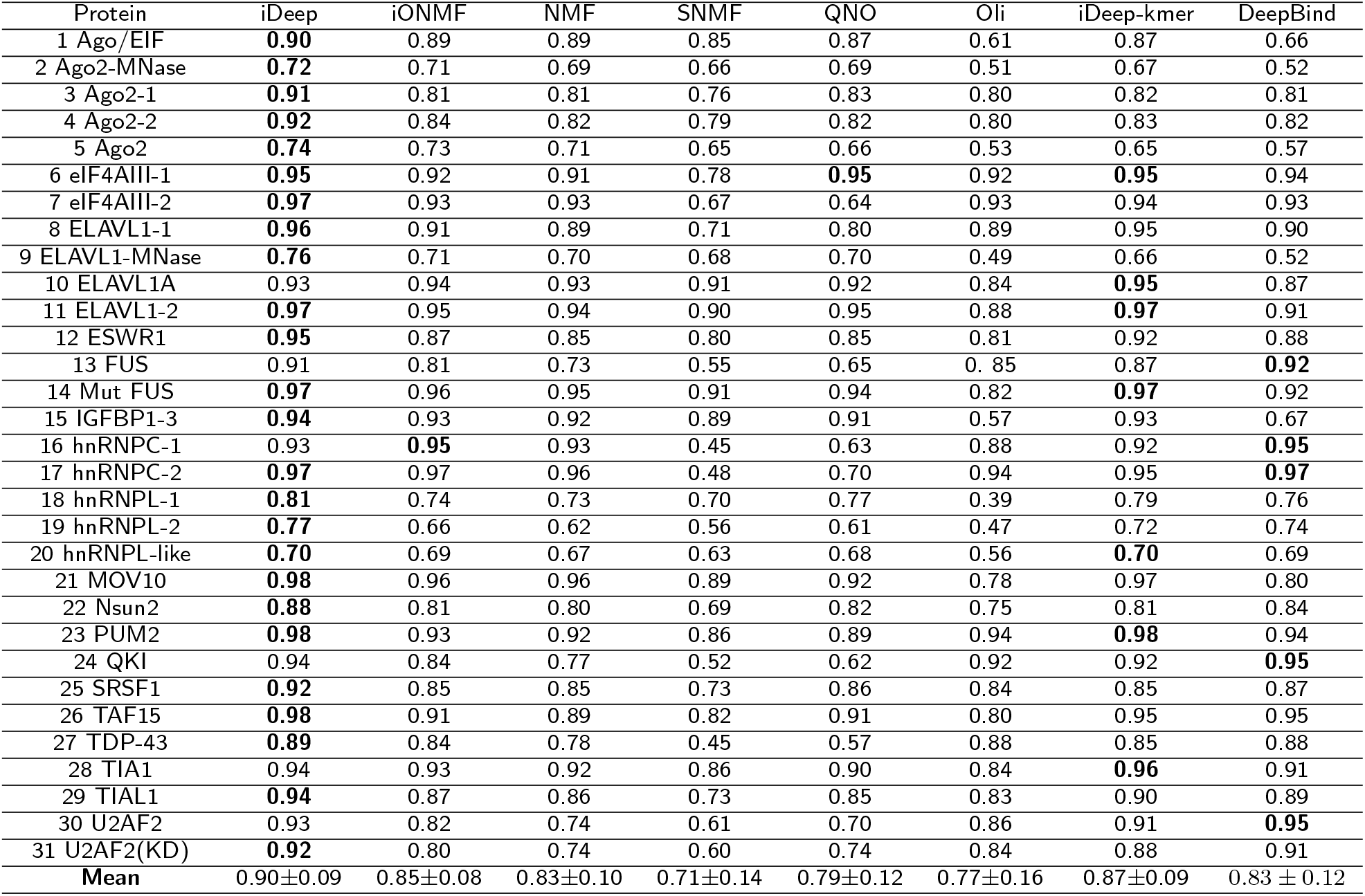
**The AUC performance comparison between iDeep and other methods on 31 experiments.** The performance of iONMF, NMF, SNMF and QNO are taken from [5]. DeepBind, Oli and iDeep-kmer perform on the same data with iDeep, and iDeep-kmer used kmer to replace CNN sequence and motif modalities in iDeep.

Overall, compared to other 6 tested methods, iDeep yields the best performance on 20 of 31 experiments and the same AUCs on other 5 experiments. And it achieves a little lower AUC only on 6 of the 31 experiments, but it still yields the AUCs above 0.90. For those experiments with AUCs below 0.90 in other six methods, iDeep’s performance is very encouraging. These results indicate that iDeep’s promising performance.

### Comparison between individual modalities

To show the advantage of integrating multiple modalities of data, we also tested the performance on individual modalities. The average AUCs of 31 experiments for region type, clip-cobinding, structure, motif and CNN sequence are 0.73 ±0.11, 0.74±0.11, 0.71±0.12, 0.71±0.08 and 0.83±0.12, respectively, indicating that individual deep networks have the ability of learning high-level features for RBP binding sites prediction. From the results, we can see that CNN sequence modality yield the best average performance with roughly 12% improvement over the second most informative region type. And CNN sequence yields higher AUC on 22 experiments due to sequence specificities of binding RNA [8], where CNN sequence can automatically learn binding motifs as feature representations for subsequent classifications. The other 4 modalities achieves similar average AUCs on all experiments without a big difference. Furthermore, we also tested the performance of DBN with only kmer modality, it yields the average AUC of 0.76±0.13 on 31 experiments, which is found much worse than CNN sequence modality.

As indicated in Fig. 3, there exists big performance differences on individual experiments for different modalities. For instance, on U2AF2 (KD) experiment, the 5 individual modalities achieve the AUC of 0.66, 0.65, 0.53, 0.72 and 0.91, respectively. The CNN sequence modality obtains AUC 0.91, outperforming other 4 modalities. While for experiment ELAVL1-MNase, they yield the AUCs of 0.67, 0.70, 0.67, 0.54, and 0.52, respectively. The CNN sequence achieves the worst AUC of 0.52 and the clip-cobinding modality has the best AUC of 0.70. The results showed that there were huge differences between different modalities on different experiments.

**Figure 3.**
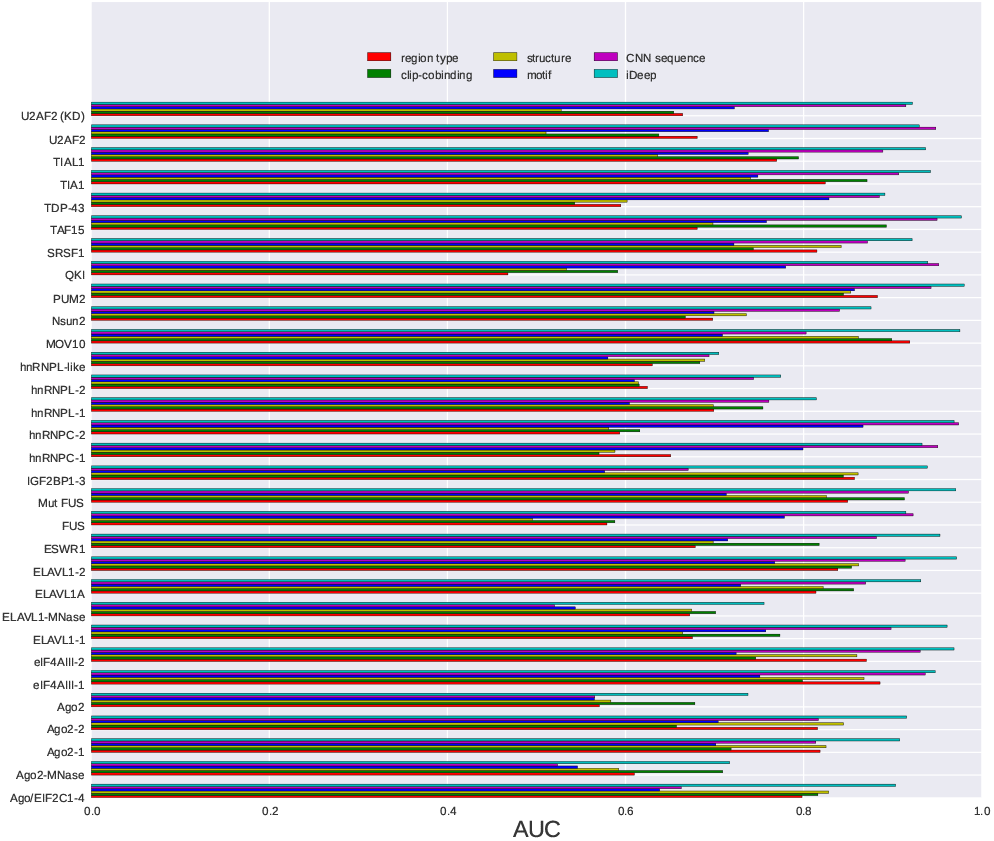
**Performance of individual modalities.** The comparison for predicting RNA-protein binding sites on 31 experiment dataset using iDeep and individual modalities.

Among the 5 Ago2 experiments, structure modality performs a little better on 3 of them. It is because that Ago2 protein requires specific RNA structure binding interfaces [41]. The motif and CNN sequence modalities perform worse than other modalities on the 5 Ago2 experiments. The reason is that Ago2 protein has a PAZ domain and a PIWI domain, but there are no related binding motifs for them in CISBP-RNA database [32], and hence deep network of motif and CNN sequence modalities cannot learn high discriminating features for predicting Ago2 binding sites on RNAs. Although motif and CNN sequence modality are not able to detect binding sites for Ago2 with high accuracy, other modalities can complement with them. The more diversity different modalities have, the more accurate the integrated method is [42]. So integrating the 5 different modalities using multimodal deep learning makes iDeep perform much better than individual modalities.

Based on the above results, we can have the following conclusions: (1) No single modality can beat others on all datasets, their performance varies on different datasets. (2) The deep network (CNN and DBN) of input modalities are able to learn high-level features with stronger discriminating ability for RBP interaction sites. (3) Integrated iDeep performs better than deep networks of individual modalities, it is because that multimodal deep learning is able to learn shared representation across multiple modalities with strong discriminating ability for RNA-protein binding sites.

### The correlations between different modalities in deep architecture

In the proposed iDeep model, we integrated 5 sources of data for an ensemble prediction. It will be interesting to see how the 5 independent modalities will complement with each other. We thus investigated the pairwise correlation between the different modalities region type, clip-cobinding, structure, motif, CNN sequence across 31 experiments. In addition, we also demonstrate the correlations between the 5 modalities and unintegrated high-dimensional k-mer modality.

We calculate the Pearson correlation coefficients (PCC) based on the AUCs of 31 experiments from individual modalities. If two modalities have high PCC, they perform similarly across all 31 experiments. As illustrated in Fig. 4, there are two obvious subgroups between the 6 modalities. The region type, clip-cobinding and structure formed the first group; kmer, motif and CNN sequence formed the other group. These results show that different modalities contain various signals, and they can complement with each other via integration in iDeep.

**Figure 4.**
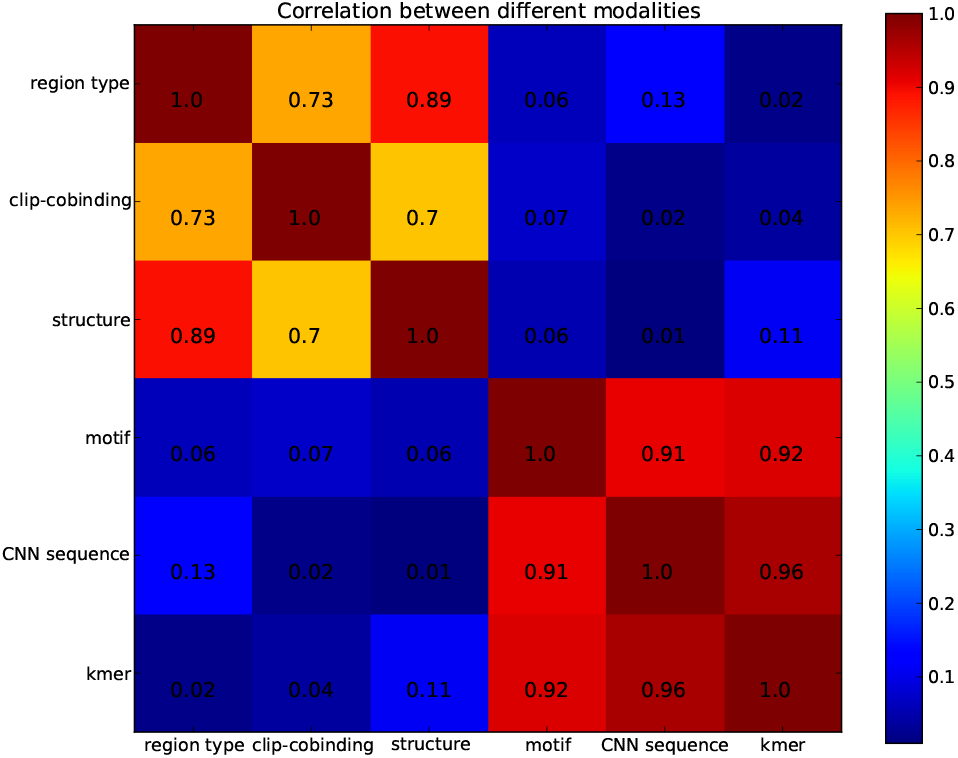
**The correlation between different modalities on 31 experiment dataset.** The pearson correlation coefficient values are calculated using the AUCs from 31 experiments for individual modalities.

The region type and structure modalities have a PCC of 0.89, showing that they are highly correlated. It is because the same region type may have similar structures, they share redundant information for predicting binding sites. CNN sequence and kmer also have very high PCC of 0.96, indicating that they are also highly related. As demonstrated in the iONMF [5], kmer modality can capture binding motifs. CNN sequence also learns motifs using CNN network [8, 19]. In addition, both of them are highly correlated to motif modality with PCCs of 0.91 and 0.92, respectively. It indicates that the high-level features learned from CNN sequences and kmer are closely related to binding motifs, which is consistent with previous findings. In summary, both the modalities try to learn binding motifs, so they share similar signals associated with motifs for RBPs across the 31 experiments. That is also the reason why we used CNN sequence instead of high-dimensional kmer in iDeep.

### The iDeep is able to discover new binding motifs

The iDeep can predict RBP binding sites on RNAs with high accuracy, however the principles behind it are still not easily interpretable. So here we further use iDeep to discover binding motifs for RBPs. In previous methods [12, 5], they focus on directly detecting nucleotide binding sites on RNAs from extracted features, but did not introduce the motifs during feature learning. Although iONMF tries to infer the binding motifs after model training, it totally depends on the input kmer sequences and defines a background distribution. In addition, it limits the learned motifs to size k, which requires optimization for searching potential motifs and the time cost exponentially increases with k.

To explore the learned motifs, we investigate the convolve filters of the convolutional layers from CNN module in iDeep, and convert them into position weight matrices (PWM), which is matched against input sequences to discover binding motifs, like DeepBind [8] and Basset [20] (Additional file 1). Then, these discovered motifs are aligned against 102 known motifs in study [32] from CISBP-RNA using the TOMTOM algorithm [43].

Using p-value <0.05, iDeep captures most of informative motifs for individual proteins. The significantly matched known motifs for individual experiments are listed in Table 2, where 14 experiments with known motifs in study [32] are included. As can be seen from Table 2 that the iDeep is able to mine known motifs for 11 of 14 experiments. For example, there are 5 known motifs (M031, M108, M112, M127, M232) in study [32] for protein ELAVL1-1, and all of them have been correctly discovered by iDeep. Fig. 5A illustrates the heatmap of learned weights of convolve filters of CNN and corresponding matched known motifs for these filters. Besides the already well-known motifs discovered by iDeep, it is able to find some novel motifs. For instance, for protein TDP-43, currently there are no verified motifs for it in CISBP-RNA database, although TDP-43 have been discovered to bind to thousands of RNAs in neuron [47]. Fig. 5B shows the hierarchical clustering of 102 new filters (motifs) for protein TDP-43 discovered by iDeep. Of them, two newly identified motif examples for protein TDP-43 are illustrated in Fig. 5C. These new motifs will provide important clues for further wet-lab verifications. All discovered motifs by iDeep are available at https://github.com/xypan1232/iDeep/tree/master/predicted_motifs.

**Table 2.**
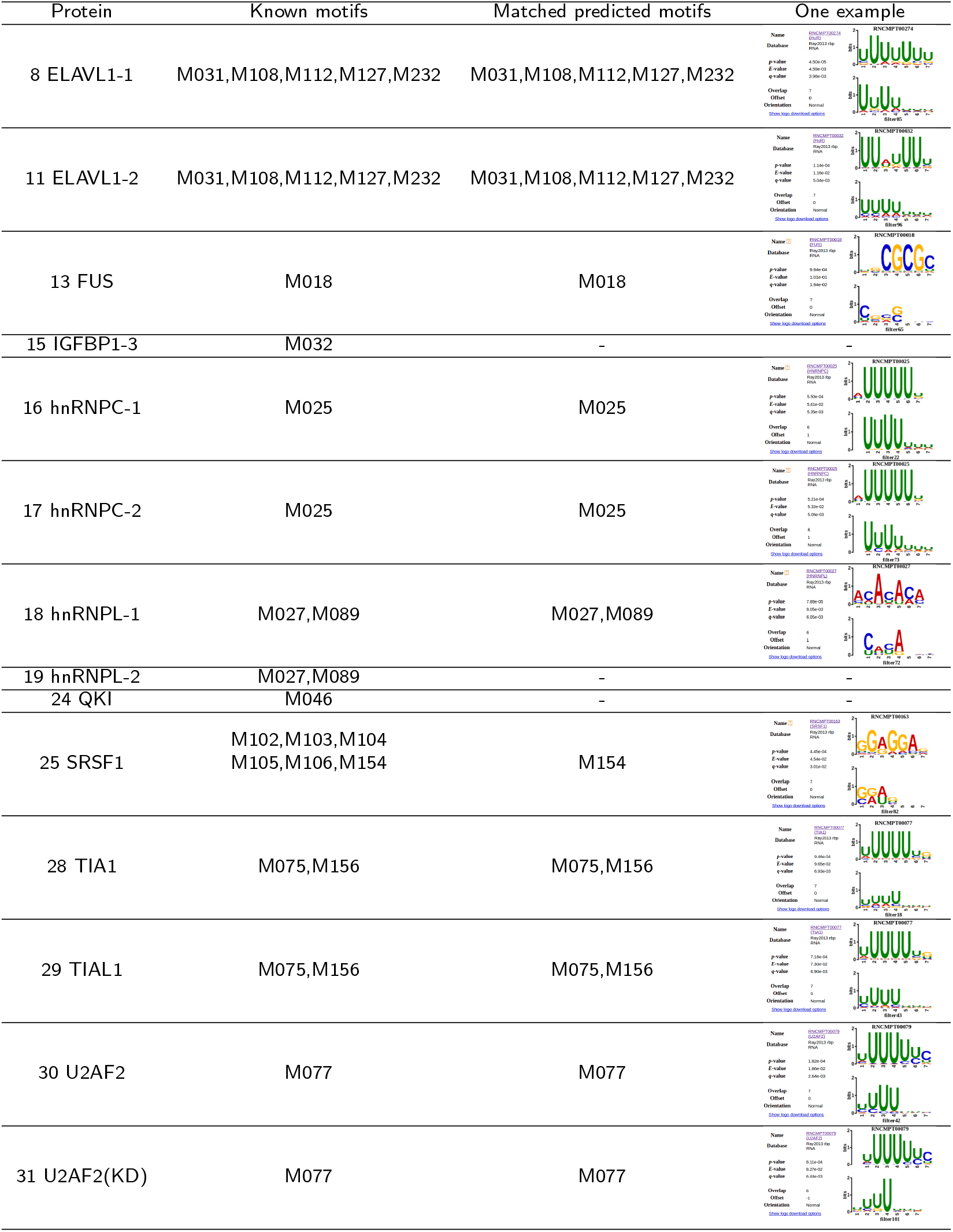
**iDeep captures known motifs in [32] from CISBP-RNA for proteins.** We only compared our predicted motifs against known motifs in study [32] and the motif name is from CISBP-RNA. If there is no motifs for this protein, then we ignore them. - means no matched motifs in our predictions.

**Figure 5.**
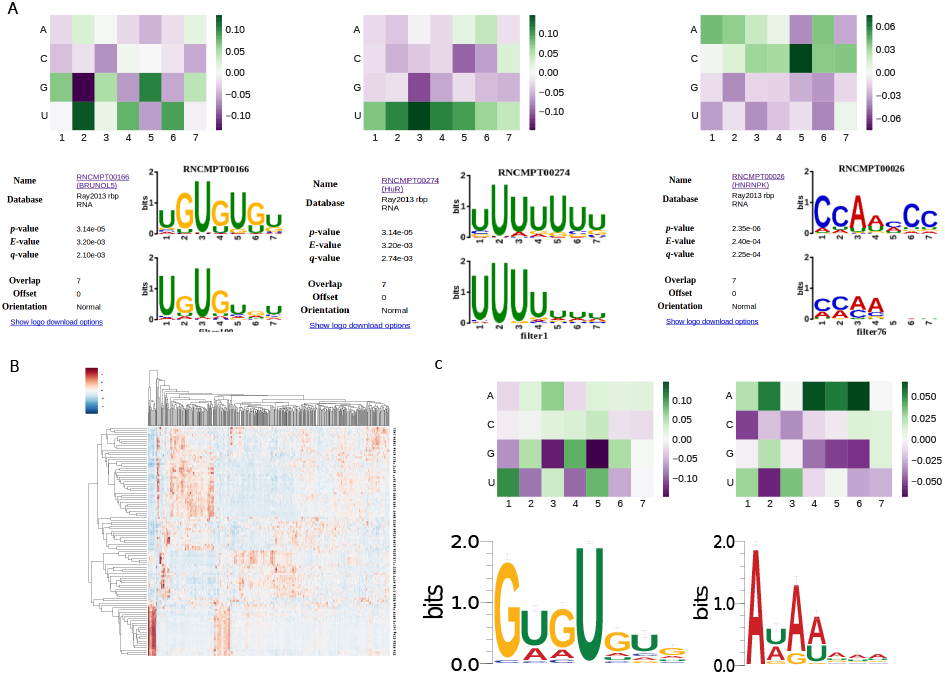
**The identified binding motifs by iDeep.** A. The heatmap of learned weights of convolve filters of CNN and corresponding matched known motifs for this filter. B. The hierarchical clustering using the cosine distance of 102 filters for protein TDP-43. C. The heatmap of learned weights of two convolve filters and corresponding motif logos for protein TDP-43, they are still not verified novel motifs detected by iDeep.

## Discussion and conclusion

In this study, we present a deep learning based hybrid framework to integrate different sources of data to predict RNA-protein binding sites on RNAs from CLIP-seq data, which yields promising performance on large-scale experiment data. The iDeep has the following advantages: (1) It trains deep neural network on individual sources of data to learn high-level representations for predicting RNA-protein interaction sites. (2) Different from other black-box machine learning based approaches, iDeep is able to discover the interpretable binding motifs, which provides better biological insights into RBPs. (3) It makes use of multimodal deep learning to extract shared features across different sources of data, with the hypothesis that no single one can overwhelm others across all datasets. Multimodal deep learning is able to better fuse them and achieve better performance on all datasets. Our proposed deep learning framework provides a powerful approach and choice for heterogeneous data integration.

In iDeep, we do not integrate high-dimensional k-mer and GO features, which possibly causes the over-fitting problem when calculating the partition functions. In addition, for other 5 integrated features in iDeep, dropout layer was applied for both CNN and DBN. It randomly sets 0s for some unit activations with certain probabilities, which can avoid over-fitting for model training [44].

In our 5 modalities integrated in iDeep, CNN sequence modality outperforms other modalities on most experiments. But for some proteins, such as Ago2, it performs worse than structure modality, indicating structure information has better informative signals for Ago2 binding sites. Currently we just use simple probabilities predicted from RNAfold [30] as the input features, which contain some noises due to the accuracy below 100%. So in future work, we will extend the CNN to structures, and design CNN to find high-level structure motifs for RBP binding sites. As done in GraphProt [6], they apply graph encoding to detect structure motifs. We can adopt similar strategy for encoding RNA structure to 6 elements (stem, multiloop, hairpin loop, internal loop, bulge and external regions), which can be fed into CNN for learning structure motifs automatically to further improve iDeep’s performance. In addition, Ago2 binding specificity is provided primarily by miRNAs [2], the expressed miRNAs in a given cell type greatly influences Ago2-RNA interactions, resulting in a much more variable and cell type-dependent binding motifs than RNA-binding proteins which bind their mRNA targets directly. Integration of miRNA expression as an additional modality will conceivably improve the accuracy of iDeep for Ago2 proteins.

The iDeep outperforms other state-of-the-art methods with the average AUC of 0.90 on 31 experiments and it can easily be used to capture binding motifs. In addition, iDeep also discovers some novel binding motifs besides those reported motifs in CISBP-RNA, we expect to verify those novel motifs by investigating whether the genes with the same predicted motifs are significantly associated with certain functions.

Despite the promising performance of iDeep, there are still promising avenues to explore the ability of deep learning. Currently we only use the standard CNNs for sequences and similar DBNs for other data modalities with only different number of hidden neurons, which should be designed specifically for different input data. Besides, more advanced network architecture could be designed according to the special characteristics of different input data. For example, DanQ designed a hybrid convolutional and recurrent neural network to predict the functions from non-coding DNA sequences [45]. It uses CNN to detect regulatory motifs from sequences, followed by bi-directional recurrent layer to capture long-term dependencies between motifs. Furthermore, instead of learning high-level features using deep learning, another study aims at automatically learning hand-designed optimization algorithms, which can exploit the structures in network architecture of interest [46]. All these studies indicate that we can further improve the structure of current iDeep to improve the performance in the future.

## Materials and Methods

In this section, we firstly introduce the CLIP-seq datasets and multiple features extracted in this study, then we design a multi data source driven multimodal deep learning framework to integrate them for predicting RNA-protein binding sites on RNAs.

### Datasets

In this study, to compare with the existing state-of-the-art methods, we used the same benchmark dataset as iONMF [5], which was downloaded from https://github.com/mstrazar/ionmf. In this dataset, the CLIP-seq data consists of 19 proteins with 31 experiments. As described in the iONMF, each nucleotide within clusters of interaction sites derived from CLIP-seq were considered as binding sites. To reduce the redundancy, the positive binding sites were further randomly sampled with the highest cDNA count and without consecutive sites on genome. Finally, from those sites with less than 15 nucleotides apart, only one site with the highest cDNA counts was selected as the positive sample. The negative sites were sampled from genes that were not identified as interacting in any of 31 experiments. In the experiments, a total 4,000 crosslinked sites are used for training purpose, 1,000 samples for model optimization and validation, and the other 1,000 samples for independent testing.

### Feature encoding

Feature encoding is critical for a statistical machine learning model. In order to integrate the merits from both the sequence and numeric features, the iDeep model makes use of 5 different groups of features, i.e., sequence, structure, clip-cobinding, region type and motif features. A scale window of [-50, 50] centering the crosslinked sites is used to generate the feature vectors, which is the same as iONMF [5].

1. **Region type**. this feature value is assigned to each position within the window using one of the 5 types (exon, intron, 5‘UTR, 3‘UTR, CDS) from Ensembl annotation [29], resulting in 101 × 5 = 505 dimensional features.
2. **clip-cobinding**. This feature represents the correlation among 31 experiments. For each experiment, the cDNA counts at each position within the window relative to the centring site was reported in the remaining 30 experiments, assign 0 for zero cDNA counts or 1 otherwise, resulting in 101 × 30=3030 dimensional features.
3. **Structure**. RNAfold [30] is used to calculate the probability of RNA secondary structure for each nucleotide within window, resulting in 101 dimensional features.
4. **Motif**. Motif scores are used for numerical representation of the RNA sequences [31]. We firstly downloaded 102 human RBP binding motifs from CISBP-RNA [32], then Cluster-Buster [33] was employed to score RNA sequences for binding sites clusters. For individual sequence, we can get a score per motif, resulting in a 102 dimensional features.
5. **CNN sequence**. The sequence is encoded into a 1000 × 4 binary matrix corresponding to the presence of A,C,G, U, which is fed into CNN to obtain high-level sequence feature.

It’s worth noting that since the iDeep model is constructed with the CNN algorithm, the 25856-D kmer and 39560-D GO features originally used in the iONMF are not used in our model. The main reasons are: 1) the GO features has been indicated of lower discriminating power than other sources of data [5] and 2) these two features are of too high dimensions, even more than the training samples, which easily leads to over-fitting and dimension disaster for neural networks. We also added two new feature encoding methods, which have not been applied in the iONMF, i.e., the sequence and motif features. Our results below will show that the new sequence feature encoding are critically important for CNNs to learn binding motifs, and the motif features based on known motifs in CISBP-RNA database are useful to correlate with functional regulatory regions in RNA sequences.

### Convolutional neural network

Convolutional neural network (CNN) is inspired by biological processes, it consists of one or more convolutional layers, followed by the max pooling layers. And it enforces a local connectivity pattern between neurons of layers to exploit spatially local structures. In this study, CNN is used to capture non-linear sequence features, e.g. motifs, and pull out some high-level features associated with RBP binding.

Here RNA sequence is one-hot representation encoded into a 101 × 4 binary matrix, whose columns correspond to A, C, G and U [8, 19]. Then the inputs are convolved with tunable patterns called filters, which are weight parameters corresponding to binding motifs and learned from RNA sequences. After convolution, a rectified linear ReLU is applied to avoid the vanishing gradient problem existing in deep learning research. Finally, a max pooling operation is used to pool adjacent positions within a small window, which can reduce the number of parameters and yield invariance to small sequence shifts.

### Deep Belief Network

Deep Belief Network (DBN) consists of multiple layers of Restricted Boltzmann machines (RBMs) [48], which learns model parameters in bottom-up style and layer-wise, but it is only able to learn abstract structure from one input source of data.

Some of our extracted input features are binary, such as region type of nucleotides. RBM is developed for binary-valued inputs, which is a graphical model with visible *υ* ∈ {0, 1} and hidden units *h* ∈ {0, 1}. Its joint distribution of hidden and visible variables are defined as follows:

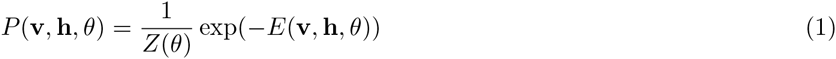

where *E*(**v**, **h**, *θ*) is defined:

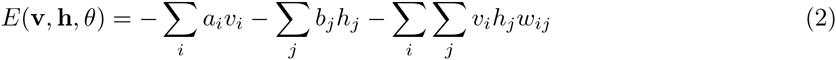

where *v_i_* and *a_i_* are binary state and bias for visible unit i, respectively. *h_j_* and *b_j_* are the binary states and bias for hidden unit j respectively.

The partition function Z is calculated by summing over hidden and visible variables, which is optimized using maximum likelihood estimation based on Contrastive Divergence algorithm [17]. Besides, we also extract structure probability features, which are real-valued inputs, and its extension Gaussian RBMs are developed for modelling real-valued inputs [34]. The parameters weight matrix and biases are updated using a gradient descent algorithm [17].

DBN is comprised of multiple RBMs, Here we take a DBN with two hidden layers as example:

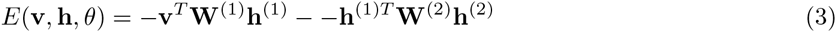

where *h*^(1)^ and *h*^(2)^ are hidden units for two hidden layers, and *W*^(1)^ and *W*^(2)^ are weight parameters for visible-to-hidden and hidden-to-hidden connection.

DBN is able to capture high-level features from individual modalities, but it cannot interactively learn unified feature representations across them.

### Multimodal deep learning for Predicting RNA-protein interaction sites

Considering the heterogeneous representations of RBP binding sites, multimodal deep learning is developed to learn shared features across different sources of data [27]. It consists of multiple layers of neural networks, which can automatically learn high-level features hidden in original features [17, 16] and achieve a huge success in different applications. In this study, we use CNNs and DBNs as the building blocks for deep learning framework shown in Fig. 1. It adds an additional layer to combine the outputs from multiple DBNs and CNNs for different inputs. During feature learning, individual DBNs and CNNs are pre-trained independently and concatenated together for final joint training using backpropagation. In each training epoch, it will automatically tune the learned parameters in respective models. After several training epochs, it learns shared representations across region type, clip-cobinding, structure, motif and CNN sequence for subsequent classification. In addition, it can also learn better features for individual modalities via backpropagation when multiple modalities exist.

We apply multimodal deep learning to integrate different sources of data to predict RNA-protein binding sites on RNAs. It first extracts different representations of different sources of data from CLIP-seq data, which are subsequently integrated using multimodal deep learning to predict RNA-protein binding sites. The flowchart is shown in Fig. 1.

In this study, we set the maximum number of epoch to 20, the batch size is 100. The neural network models are optimized using RMSprop algorithm [35] to learn all model parameters, including those convolution filters of CNNs. Validation dataset is evaluated to monitor the convergence during each epoch of the training process, so the training process can be stopped early.

The iDeep is implemented in python using keras 1.0.4 library https://github.com/fchollet/keras. The model architecture consists of hybrid CNNs and DBNs for individual inputs and additional layer for merging them.

For sequence modality, its one-hot encoding is fed into CNN to learn high-level motif features. The parameter nb filter (number of motifs) is 102 and the filter length (motif width) is 7, which agrees with the significantly verified RBP binding motifs in CISBP-RNA database [32].

The architecture of DBN for input modalities clip-cobinding, Structure, Region type and Motif consists of fully connected layer and dropout layers (Additional file 1). In iDeep, for each DBN from individual modalities, we configure different number of hidden units for two Fully connected layer (FCL) listed in Table 3, and the dropout probability for each dropout layer is 0.5. To evaluate the performance of predicting RBP binding sites, we use Receiver Operating Characteristic(ROC) curve and calculate the area under the ROC curve (AUC).

**Table 3.**
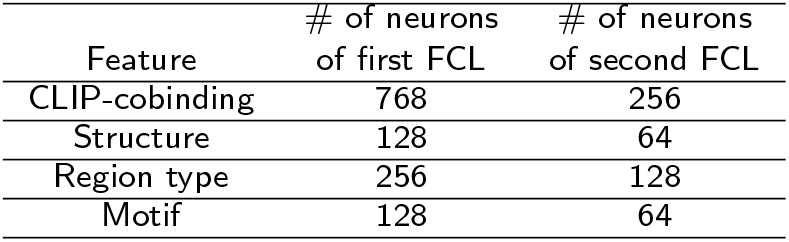
The number of neurons used in Fully connected layer (FCL) for each DBN.

### Baseline methods

There are many computational methods developed for predicting RNA-protein binding sites. such as iONMF, Oli, DeepBind, GraphProt and RNAContext. As indicated in [5], iONMF performs a little better than GraphProt, and much better than RNAContext. In [12]), Oli with only tetranucleotide frequency features yield better performance than its variant OliMoSS for predicting RBP binding sites. So in this study, we compared iDeep with other state-of-the-art iONMF, DeepBind and Oli. iONMF integrates multiple data using orthogonality-regularized nonnegative matrix factorization, it discovers the hidden modules from non-overlapping features for RNA-protein interactions. Oli applied linear SVC to classify protein-RNA binding sites based on their extracted tetranucleotide frequency features. To compare with Oli fairly, grid-search was used to select the best parameter for linear SVC of Oli in individual experiments, and the implementation from scikit-learn package was used in this study [36]. For DeepBind, it only uses CNN from sequences to predict RBP binding sites.

## Abbreviations

RBPs: RNA binding proteins
RISC: RNA-induced silencing complex
CNN: convolutional neural network
DBN: Deep belief network
FCL: Fully connected layer
RF: Random Forest
SVM: Support vector machine
ROC: Receiver Operating Characteristic
GO: Gene Ontology
AUC: the area under the ROC curve
PCC: Pearson correlation coefficient
PWM: position weight matrices

## Declarations

Ethics approval and consent to participate

Not applicable

### Consent for publication

Not applicable

### Availability of data and code

The datasets and python code supporting the findings of this study are available at https://github.com/xypan1232/iDeep or http://www.csbio.sjtu.edu.cn/bioinf/iDeep.

### Author’s contributions

XP and HBS designed the study and drafted the manuscript, XP did the bioinformatics analysis. All authors read and approved the final manuscript.

---

Additional file 1 — Supplementary text and Table

Some details of iDeep. The principles about how to identify binding motifs by iDeep, the architecture of deep belief network and the discovered number of known motifs in CISBP-RNA.

